# Antimicrobial susceptibility patterns of anaerobic bacteria in Victoria, Australia

**DOI:** 10.1101/308023

**Authors:** Jasmine C. Teng, L Linda Dreyer

## Abstract

Mortality associated with anaerobic infections approximates 20%. Resistance of anaerobic bacteria to commonly used antimicrobials has been increasingly reported. The aim of this study was to describe antimicrobial susceptibility patterns of anaerobic bacteria isolated from clinical samples using a gradient diffusion method, E test (bioMérieux), in Victoria, Australia. Metronidazole, meropenem and amoxycillin-clavulanate were found to be active against almost all isolates tested. Most Gram positive anaerobic cocci (GPAC), except *Peptostreptococcus anaerobius* (64.6% penicillin-susceptible), remained susceptible to penicillin. All *Clostridium perfringens* isolates tested were penicillin, metronidazole and meropenem susceptible. Of *B. fragilis* isolates tested, 5% and 0.83% were meropenem and metronidazole non-susceptible, respectively. Clindamycin susceptibility in anaerobes other than the GPAC is approximately 75% and therefore should not be used as empirical treatment in the absence of susceptibility testing. Considering the global trend of antibiotic resistance among anaerobic bacteria, routine susceptibility testing of anaerobic bacteria, particularly when isolated from critical sites, as well as surveillance of local resistance trends is strongly encouraged. Gradient diffusion MIC determination of anaerobic bacteria is feasible in a clinical diagnostic laboratory and should be more widely utilised.

## Introduction

Anaerobic infections cause significant morbidity and mortality and various clinical studies have demonstrated adverse survival outcomes in patients due to inappropriate therapy. Furthermore, anaerobic resistance to commonly used antimicrobial agents has increasingly been reported.(1, 2)

Despite this, routine antimicrobial susceptibility testing of clinical anaerobic isolates remains a contentious issue.(3) This is in part due to difficulties associated with identification, purification and manipulation of anaerobes. In the past decade, the introduction of matrix-assisted light desorption ionization-time of flight (Maldi-TOF) mass spectrometry in most diagnostic microbiology laboratories has greatly enhanced the ability for microbial identification in a time and cost-efficient manner. Multiple studies have corroborated the accuracy and reliability of anaerobic bacteria identification by MALDI-TOF.(4–7) At present, there is no ISO standard reference method for susceptibility testing of anaerobic bacteria. Procedural guidelines have been published by the Clinical and Laboratory Standards Institute (CLSI), (8) European Committee on Antimicrobial Susceptibility Testing (EUCAST),(9) and Calibration, Dichotomous Susceptibility (CDS).(10) CLSI recommends minimum inhibitory concentration (MIC) determination by broth microdilution for Bacteroides fragilis group and agar dilution for all anaerobes,(8) whereas the EUCAST recommends testing with an MIC method, and reference to the manufacturer’s instructions of a commercial product.(9) Clinical MIC breakpoints for main classes of anaerobic antimicrobials are provided by each committee respectively; these differ and should be interpreted with care. The CDS recommends disc susceptibility testing method for anaerobes with interpretive annular radius cutoffs.(10)

Both broth microdilution and agar dilution methods for anaerobes are time-consuming, require expertise and are not practical to be implemented in a routine diagnostic laboratory. In the past, susceptibility testing by disc diffusion has not been recommended due to suboptimal correlation and reproducibility.(11) A recent correlation study by Nagy et al, demonstrated good agreement between zone diameter and MIC for Bacteroides fragilis group of bacteria using EUCAST rules and breakpoints,(12) although further validation is needed. MIC determination by gradient diffusion has shown reasonable correlation with broth microdilution and agar dilution methods.(13–17) Gradient diffusion MIC is easy to perform and readily implemented using commercially available products, E test (bioMérieux) and MIC Evaluator (M.I.C.E., Thermo Fisher Scientific) strips.

In Australia, antimicrobial susceptibility testing of anaerobes is done sporadically and treatment of anaerobic infections is largely empirical. As such, antimicrobial susceptibility trends of anaerobic bacteria over time is largely unknown.

The aim of this study was to describe the antimicrobial susceptibility patterns of anaerobic bacteria from clinical samples in a private clinical microbiology laboratory in Victoria, Australia.

## Materials and methods

### Isolates

Anaerobic bacteria isolated from clinical specimens were collected from January 2015 to January 2018 from both community and hospital samples from Victoria. Clinical sources included blood culture, swabs of skin/superficial sites, deep sites/abscesses, wounds (not otherwise specified) and genital swabs. Pure anaerobic bacterial isolates were obtained from blood culture and sterile sites; the presence of anaerobes from polymicrobial or non-sterile sites is indicated by a zone of inhibition around a metronidazole (5μg) disc as per laboratory protocol. Using the Bruker matrix-assisted laser desorption/ionization-time of flight (MALDI-TOFMS) instrument, anaerobic bacteria were identified to species and genus level based on log(score) ≥2.0 and log(score) 1.7-2.0, respectively.

### Susceptibility testing

#### Gradient diffusion MIC determination

A 1 MacFarland standard suspension of a 48hr growth culture was made and inoculated with a swab onto Brucella Agar supplemented with blood (5%), 5mg/L haemin and 1mg/L vitamin K (Oxoid PP2459). E-test strips (bioMérieux) were then applied and the plates incubated at 35^°^C, under anaerobic conditions using an atmosphere generation system (AnaeroGen, Oxoid, AN0035A). *Bacteroides fragilis* ATCC25285 was tested against each new lot number of Etest strips as quality control, as per the manufacturer’s instructions. Controls for anaerobiasis included organism controls, *Bacteroides fragilis* ATCC25285 and *Pseudomonas aeruginosa* ATCC25668, and a chemical resazurin redox indicator (Anaerobic Indicator, Oxoid, BR0055B). Each isolate was tested against the antibiotics benzylpenicillin, amoxycillin-clavulanate, clindamycin, metronidazole and meropenem. MIC was read at 100% growth inhibition after 24 and 48hours of incubation. Results obtained at 48hours were considered final. The MIC values were interpreted according to both the CLSI(8) and EUCAST(9) clinical breakpoints. The amoxycillin-clavulanate E test strips contained amoxycillin and clavulanic acid in a 2:1 ratio, therefore only CLSI breakpoints were applied for interpretation. Slower growing anaerobic bacteria and those which could not be identified reliably by Maldi-TOF MS were excluded from this study.

## Results/Discussion

Four hundred and sixteen anaerobic isolates were collected during the study period. Clinical sources for these bacteria included blood culture(n=68), swabs of skin and superficial sites(n=77), deep collection/abscesses (n=37), wounds (not otherwise specified) (n=216), and genital swabs(n=19). The anaerobic bacteria collected and tested are shown in Table 1. *Bacteroides fragilis* (n=120) and *Peptostreptococcus anaerobius* (n=79) were the most commonly isolated anaerobes. The MIC range, MIC_50_, MIC_90_ and percentage susceptibility according to CLSI and EUCAST breakpoints for the Gram negative and Gram positive anaerobes are shown in Tables 2.

**Table 1:** List of anaerobic bacteria isolated between January 2015-January 2018

| **<u>Anaerobic bacteria</u>** | **<u>No. of isolates</u>** |
| --- | --- |
| *Actinomyces spp.* |  |
| *A.europeus* | 2 |
| *A.turiciensis* | 1 |
| *Anaerococcus spp.* |  |
| *A.hydrogenalis* | 6 |
| *A.murdochii* | 2 |
| *A.octavius* | 1 |
| *A.vaginalis* | 10 |
| *Bacteroides spp.* |  |
| *B. caccae* | 3 |
| *B. cellulosilyticus* | 1 |
| *B. faecis* | 3 |
| *B. fragilis* | 83 |
| *B. ovatus* | 7 |
| *B. pyogenes* | 6 |
| *B. stercoris* | 2 |
| *B. thetaiotaomicron* | 16 |
| *B. uniformis* | 3 |
| *B. vulgatus* | 5 |
| *Bifidobacterium dentium* | 1 |
| *Clostridium spp.* |  |
| *C. paraputrificum* | 2 |
| *C. perfringens* | 30 |
| *C. ramosum* | 2 |
| *C. septicum* | 4 |
| *C. sporogenes* | 3 |
| *C. tertium* | 2 |
| *Disulfovibrio desulfuricans* | 1 |
| *Eggerthia catenaformis* | 1 |
| *Finegoldia magna* | 47 |
| *Murdochiella asaccharolytica* | 1 |
| *Odoribacter splanchnicus* | 1 |
| *Parvimonas micra* | 2 |
| *Peptoniphilus harei* | 38 |
| *Peptostreptococcus anaerobius* | 79 |
| *Prevotella sp* |  |
| *P. bivia* | 24 |
| *P. buccae* | 1 |
| *P. disiens* | 4 |
| *P. nanceiensis* | 1 |
| *P. timonensis* | 1 |
| *Propionebacterium spp.* | 17 |
| *Ruminococcus gnavus* | 1 |
| *Staphylococcus saccharolyticus* | 2 |
| Total | 416 |

**Table 2:**
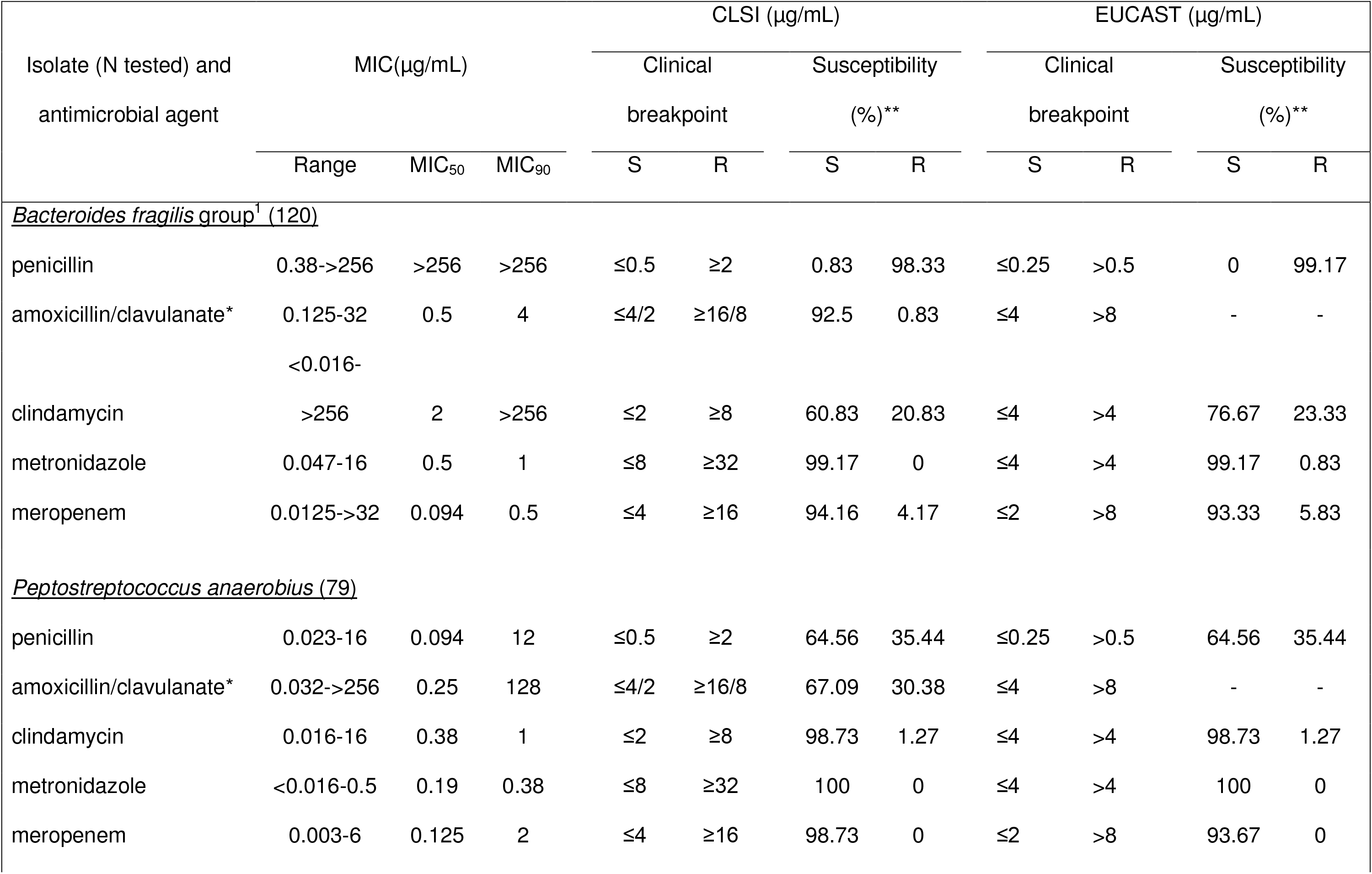
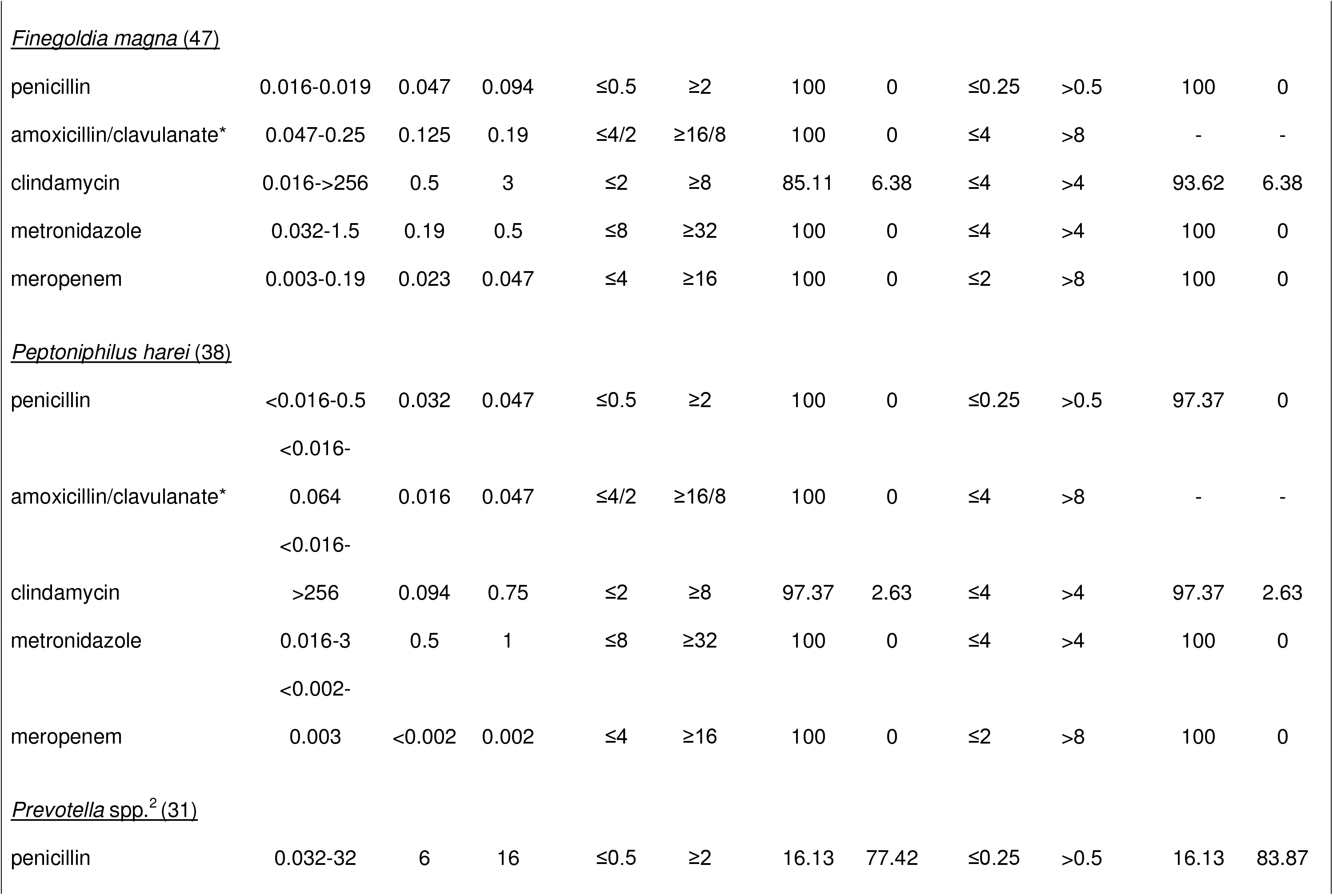
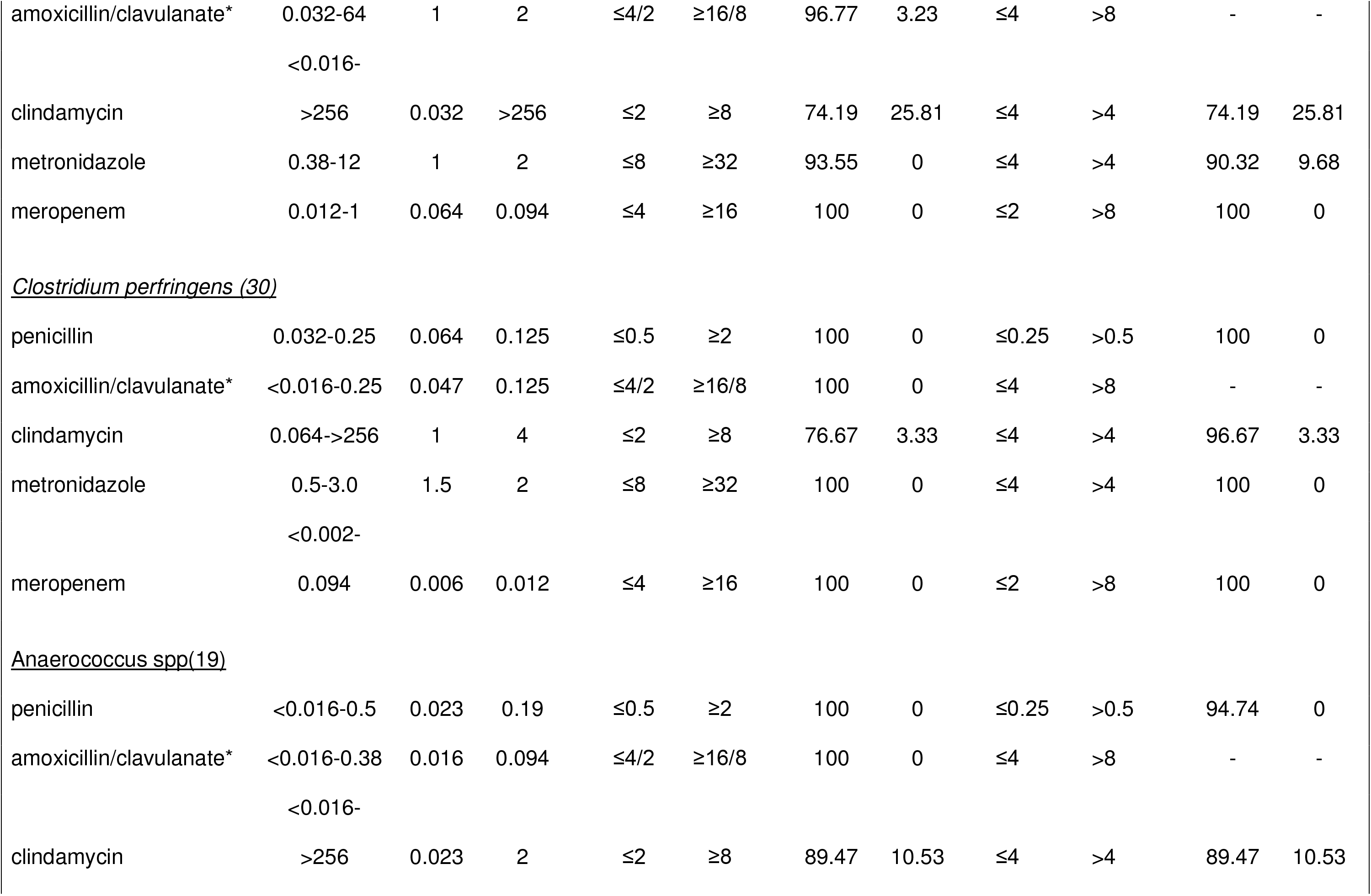
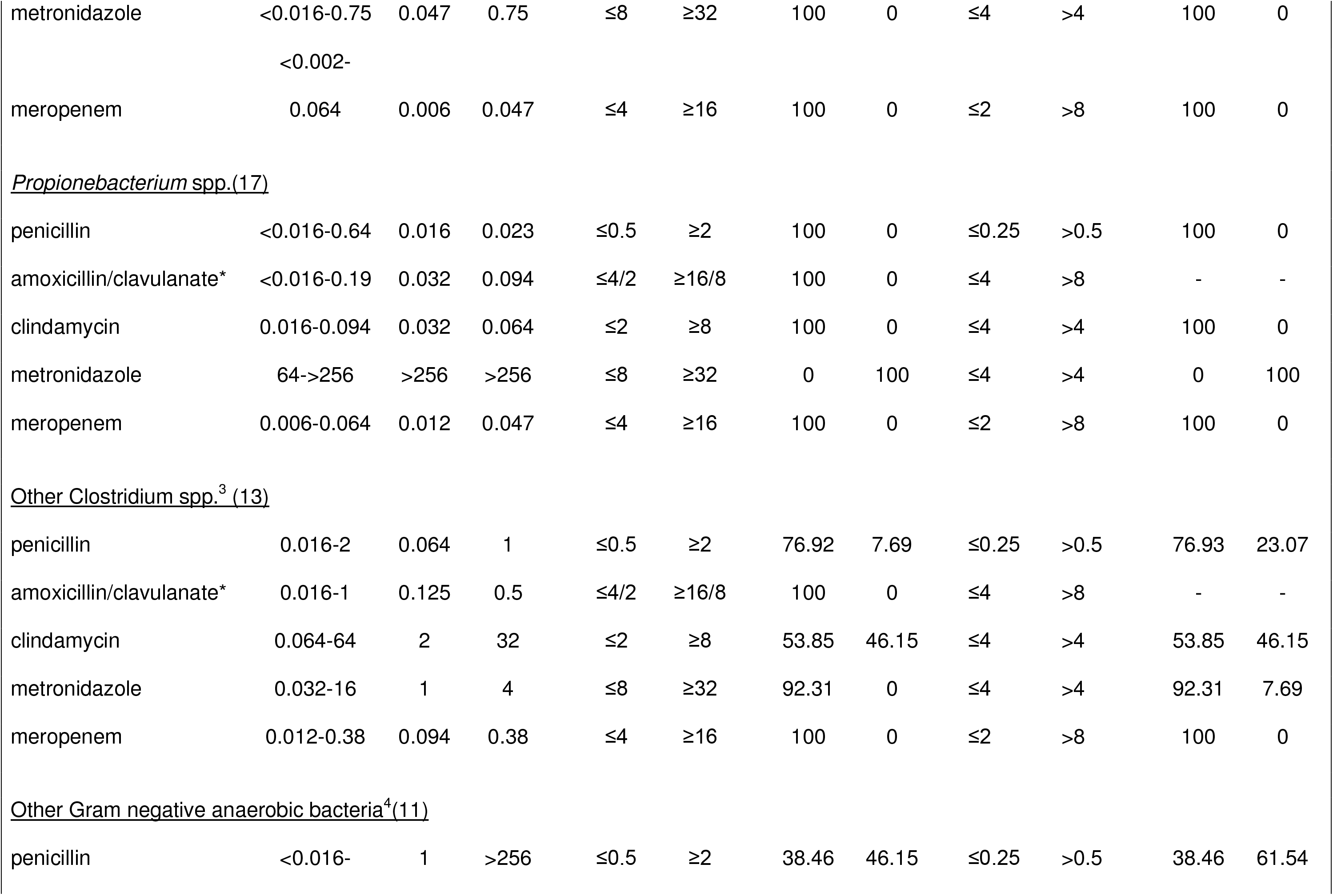
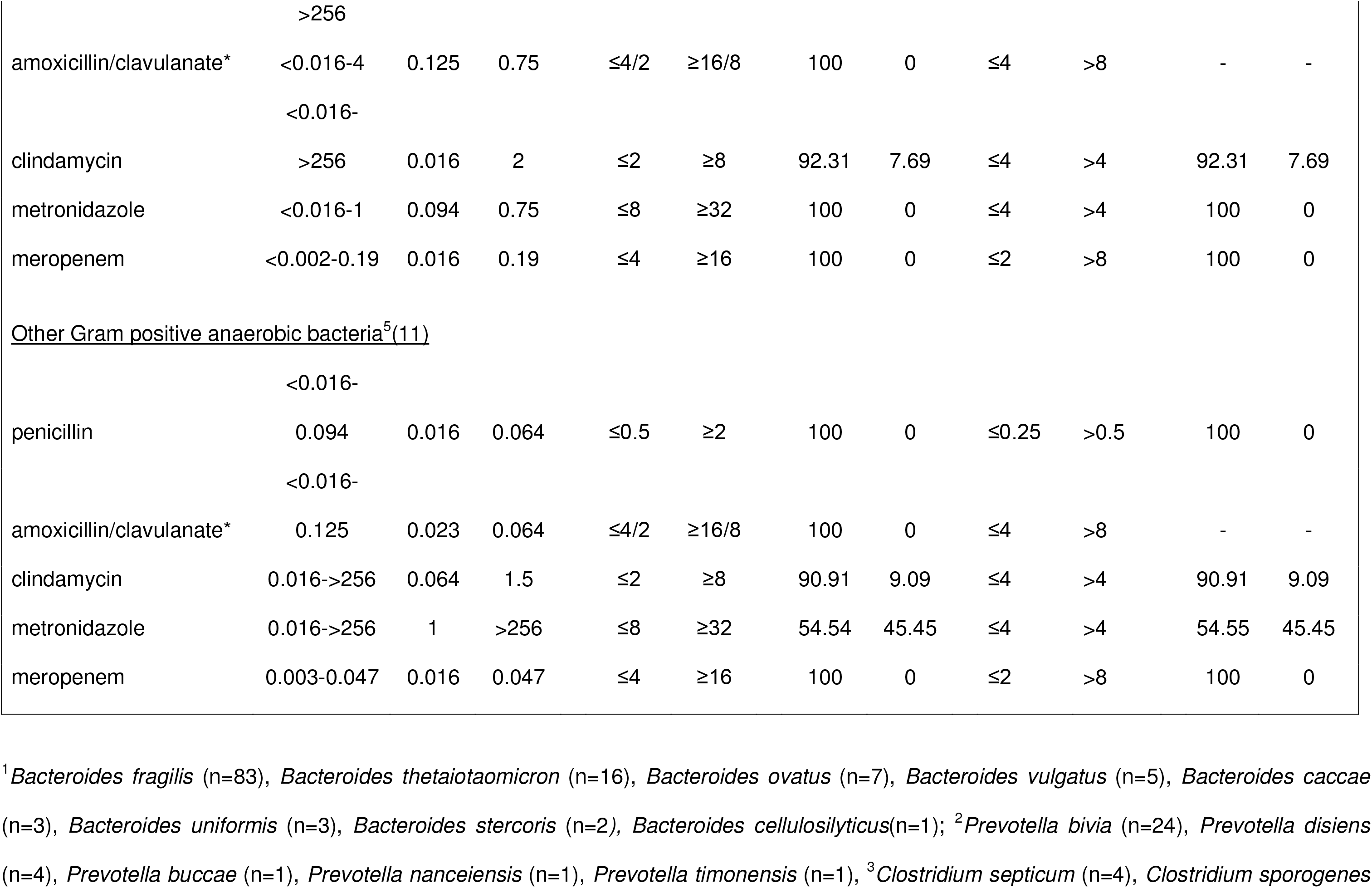
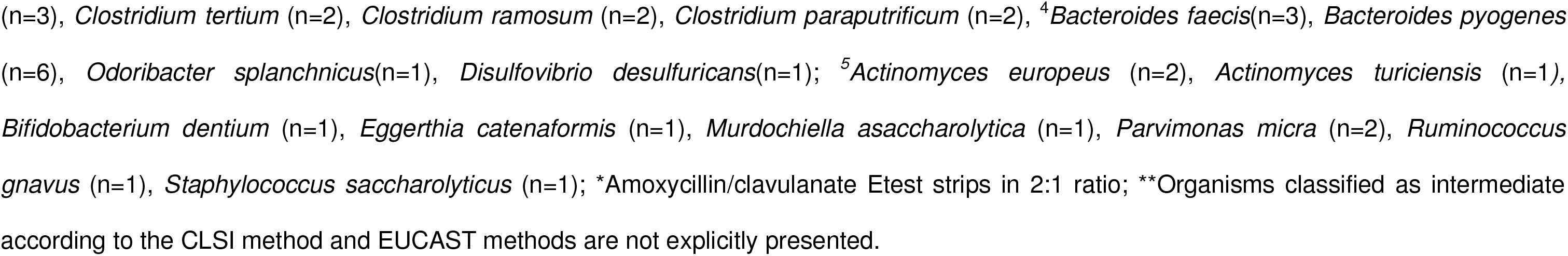
Susceptibility patterns of anaerobic Gram-negative isolates to penicillin, clindamycin, metronidazole and meropenem.

Consistent with known data, all isolates in the *Bacteroides fragilis* group and most *Prevotella* isolates (83.7%) were penicillin non-susceptible. 99.1% and 94% *B. fragilis* isolates were susceptible to metronidazole and meropenem, respectively. Most Gram positive anaerobic cocci (GPAC), except *Peptostreptococcus anaerobius* (64.6% penicillin-susceptible), remained susceptible to penicillin. 100% *Propionebacterium* sp. tested (n=17) were metronidazole-resistant and susceptible to penicillin and clindamycin. All *Clostridium perfringens* isolates tested were penicillin, metronidazole and meropenem susceptible. Clindamycin susceptibility varied across all groups of anaerobic bacteria. When both CLSI and EUCAST MIC breakpoints were applied, the overall categorical agreement for penicillin, clindamycin, metronidazole and meropenem were 97.6%, 92.3%,99.0% and 98.3%, respectively.

Anaerobic bacteria form part of the normal human indigenous microflora.(18) Research into the remarkable diversity of the human microbiome, including anaerobic bacteria in health and disease states has flourished in recent years, with particular emphasis on gut microbiome.(18, 19) Anaerobic bacteria are also opportunistic pathogens, causing bacteraemia and sepsis(20), necrotizing skin infections,(21) and rarely endocarditis.(22) The mortality rate associated with anaerobic bacteraemia has been reported to be 1-19%.(20, 23) Publications on antimicrobial resistance trends among anaerobic bacteria in North America(24, 25), Europe(2, 26, 27), and Australasia(28, 29) are prolific. Overall there has been increase in cfiA gene encoded chromosomal zinc metallo-β-lactamase enzyme mediated carbapenem resistance and resistance to clindamycin and metronidazole. According to our study, metronidazole, meropenem and amoxycillin-clavulanate were found to be active against almost all isolates tested, making them ideal agents for empirical therapy. Of the 120 *B. fragilis* isolates tested, 6 (5%) and 1 (0.83%) were meropenem and metronidazole non-susceptible, respectively. Clindamycin susceptibility in anaerobes other than the GPAC was approximately 75% and therefore should not be used as empirical treatment in the absence of susceptibility testing. Of note, 100% of *Propionebacterium* spp. was found to be metronidazole-resistant; this resistance profile may reliably be used as a supplementary for organism identification; conversely, in cases of *Propionebacterium* spp. postoperative shoulder joint or central nervous system shunt infections, metronidazole should not be used as a therapeutic agent.

The use of a metronidazole (5μg) disc for screening and detection of anaerobes in polymicrobial and non-sterile samples has been part of laboratory practice for decades. While this method is cheap and simple, it biases towards isolation of susceptible strains and will inherently miss metronidazole resistant strains. Thus, the proportion of metronidazole-resistant anaerobic bacterial isolates from non-sterile sites in this study is likely underestimated.

Gradient diffusion MIC determination using commercially available products E tests (bioMérieux) and MIC Evaluator (M.I.C.E., Thermo Fisher Scientific) strips are easy to setup and use without the need of specialized equipment. According to our experience, MIC determination of the commonly encountered anaerobes at 100% growth inhibition read at 48 hours was straightforward, with minimal interobserver variability. Therefore, this method is uniquely placed for susceptibility testing of anaerobic bacteria in routine clinical microbiology practice, and should be more commonly utilized. In this study, carbapenem and metronidazole resistant isolates by E test was not confirmed with agar dilution or molecular detection of resistance genes.

This study provides MIC data on the current local resistance patterns of commonly encountered anaerobes in Australia by gradient diffusion. Considering the global trend of antibiotic resistance among aerobic bacteria, routine susceptibility testing of anaerobic bacteria, particularly isolates from critical sites, as well as surveillance of local resistance trends is strongly encouraged.

## Acknowledgements

Contribution statement: LD and JT jointly conceived the study idea and design. JT performed susceptibility testing on all isolates. JT and LD contributed to data analysis, manuscript writing and review of manuscript. The authors would like to acknowledge all microbiology scientists at Australian Clinical Labs for assistance in collection of isolates.

## References

1. Hastey CJ, Boyd H, Schuetz AN, Anderson K, Citron DM, Dzink-Fox J, Hackel M, Hecht DW, Jacobus NV, Jenkins SG, Karlsson M, Knapp CC, Koeth LM, Wexler H, Roe-Carpenter DE, From the Ad Hoc Working Group on Antimicrobial Susceptibility Testing of Anaerobic Bacteria of C. 2016. Changes in the antibiotic susceptibility of anaerobic bacteria from 2007-2009 to 2010-2012 based on the CLSI methodology. Anaerobe doi:10.1016/j.anaerobe.2016.07.003.

2. Boyanova L, Kolarov R, Mitov I. 2015. Recent evolution of antibiotic resistance in the anaerobes as compared to previous decades. Anaerobe 31:4–10.

3. Nagy E, Schuetz A. 2015. Is there a need for the antibiotic susceptibility testing of anaerobic bacteria? Anaerobe 31:2–3.

4. Veloo AC, Welling GW, Degener JE. 2011. The identification of anaerobic bacteria using MALDI-TOF MS. Anaerobe 17:211–212.

5. Justesen US, Holm A, Knudsen E, Andersen LB, Jensen TG, Kemp M, Skov MN, Gahrn-Hansen B, Moller JK. 2011. Species identification of clinical isolates of anaerobic bacteria: a comparison of two matrix-assisted laser desorption ionization-time of flight mass spectrometry systems. J Clin Microbiol 49:4314–4318.

6. Barba MJ, Fernandez A, Oviano M, Fernandez B, Velasco D, Bou G. 2014. Evaluation of MALDI-TOF mass spectrometry for identification of anaerobic bacteria. Anaerobe 30:126–128.

7. Hsu YM, Burnham CA. 2014. MALDI-TOF MS identification of anaerobic bacteria: assessment of pre-analytical variables and specimen preparation techniques. Diagn Microbiol Infect Dis 79:144–148.

8. Anonymous. 2012. Clinical Laboratory Standards Institute, Methods for Antimicrobial Susceptibility Testing of Anaerobic Bacteria. Approved Standard CLSI Publication Number M11-a8.

9. Anonymous. 2017. The European Committee on Antimicrobial Susceptibility Testing, Breakpoint Tables for Interpretation of MICs and Zone Diameters. Version 7, 2017.

10. Anonymous. 2016. Antibiotic Susceptibility Testing By The CDS Method. A Manual For Medical And Veterinary Laboratories. Eighth Edition.

11. Schuetz AN. 2014. Antimicrobial resistance and susceptibility testing of anaerobic bacteria. Clin Infect Dis 59:698–705.

12. Nagy E, Justesen US, Eitel Z, Urban E, Infection ESGoA. 2015. Development of EUCAST disk diffusion method for susceptibility testing of the Bacteroides fragilis group isolates. Anaerobe 31:65–71.

13. Rennie RP, Turnbull L, Brosnikoff C, Cloke J. 2012. First comprehensive evaluation of the M.I.C. evaluator device compared to Etest and CLSI reference dilution methods for antimicrobial susceptibility testing of clinical strains of anaerobes and other fastidious bacterial species. J Clin Microbiol 50:1153–1157.

14. Rosenblatt JE, Gustafson DR. 1995. Evaluation of the Etest for susceptibility testing of anaerobic bacteria. Diagn Microbiol Infect Dis 22:279–284.

15. Croco JL, Erwin ME, Jennings JM, Putnam LR, Jones RN. 1995. Evaluation of the Etest for determinations of antimicrobial spectrum and potency against anaerobes associated with bacterial vaginosis and peritonitis. Clin Infect Dis 20 Suppl 2:S339–341.

16. Citron DM, Ostovari MI, Karlsson A, Goldstein EJ. 1991. Evaluation of the E test for susceptibility testing of anaerobic bacteria. J Clin Microbiol 29:2197–2203.

17. Wust J, Hardegger U. 1992. Comparison of the E test and a reference agar dilution method for susceptibility testing of anaerobic bacteria. Eur J Clin Microbiol Infect Dis 11:1169–1173.

18. Hentges DJ. 1993. The anaerobic microflora of the human body. Clin Infect Dis 16 Suppl 4:S175–180.

19. Lloyd-Price J, Abu-Ali G, Huttenhower C. 2016. The healthy human microbiome. Genome Med 8:51.

20. Brook I. 2010. The role of anaerobic bacteria in bacteremia. Anaerobe 16:183–189.

21. Zhao-Fleming H, Dissanaike S, Rumbaugh K. 2017. Are anaerobes a major, underappreciated cause of necrotizing infections? Anaerobe 45:65–70.

22. Kestler M, Munoz P, Marin M, Goenaga MA, Idigoras Viedma P, de Alarcon A, Lepe JA, Sousa Regueiro D, Bravo-Ferrer JM, Pajaron M, Costas C, Garcia-Lopez MV, Hidalgo-Tenorio C, Moreno M, Bouza E, Spanish Collaboration on E. 2017. Endocarditis caused by anaerobic bacteria. Anaerobe 47:33–38.

23. Goldstein EJ. 1996. Anaerobic bacteremia. Clin Infect Dis 23 Suppl 1:S97–101.

24. Snydman DR, Jacobus NV, McDermott LA, Golan Y, Goldstein EJ, Harrell L, Jenkins S, Newton D, Pierson C, Rosenblatt J, Venezia R, Gorbach SL, Queenan AM, Hecht DW. 2011. Update on resistance of Bacteroides fragilis group and related species with special attention to carbapenems 2006-2009. Anaerobe 17:147–151.

25. Snydman DR, Jacobus NV, McDermott LA, Goldstein EJ, Harrell L, Jenkins SG, Newton D, Patel R, Hecht DW. 2017. Trends in antimicrobial resistance among Bacteroides species and Parabacteroides species in the United States from 2010-2012 with comparison to 2008-2009. Anaerobe 43:21–26.

26. Ferlov-Schwensen SA, Sydenham TV, Hansen KCM, Hoegh SV, Justesen US. 2017. Prevalence of antimicrobial resistance and the cfiA resistance gene in Danish Bacteroides fragilis group isolates since 1973. Int J Antimicrob Agents 50:552–556.

27. Wybo I, Van den Bossche D, Soetens O, Vekens E, Vandoorslaer K, Claeys G, Glupczynski Y, Ieven M, Melin P, Nonhoff C, Rodriguez-Villalobos H, Verhaegen J, Pierard D. 2014. Fourth Belgian multicentre survey of antibiotic susceptibility of anaerobic bacteria. J Antimicrob Chemother 69:155–161.

28. Roberts SA, Shore KP, Paviour SD, Holland D, Morris AJ. 2006. Antimicrobial susceptibility of anaerobic bacteria in New Zealand: 1999-2003. J Antimicrob Chemother 57:992–998.

29. Chen SC, Gottlieb T, Palmer JM, Morris G, Gilbert GL. 1992. Antimicrobial susceptibility of anaerobic bacteria in Australia. J Antimicrob Chemother 30:811–820.

